# 2FAST2Q: A general-purpose sequence search and counting program for FASTQ files

**DOI:** 10.1101/2021.12.17.473121

**Authors:** Afonso M. Bravo, Athanasios Typas, Jan-Willem Veening

## Abstract

The increasingly widespread use of next generation sequencing protocols has brought the need for the development of user-friendly raw data processing tools. Here, we present 2FAST2Q, a versatile and intuitive standalone program capable of extracting and counting feature occurrences in FASTQ files. 2FAST2Q can be used in any experimental setup that requires feature extraction from raw reads, being able to quickly handle mismatch alignments, nucleotide wise Phred score filtering, custom read trimming, and sequence searching within a single program. Using published CRISPRi datasets in which *Escherichia coli* and *Mycobacterium tuberculosis* gene essentiality, as well as host-cell sensitivity towards SARS-CoV2 infectivity were tested, we demonstrate that 2FAST2Q efficiently recapitulates the output in read counts per provided feature as with traditional pipelines. Moreover, we show how different FASTQ read filtering parameters impact downstream analysis, and suggest a default usage protocol. 2FAST2Q has a familiar user interface and uses a custom sequence mismatch search algorithm, taking advantage of Python’s numba module JIT runtime speeds. It is thus easier to use and faster than currently available tools, efficiently processing large CRISPRi-Seq or random-barcode sequencing datasets on any up-to-date laptop. 2FAST2Q is available as an executable file for all current operating systems without installation and as a Python3 module on the PyPI repository (available at https://veeninglab.com/2fast2q). We expect that 2FAST2Q will not only be useful for people working in microbiology but also for other fields in which amplicon sequencing data is generated.

## Introduction

Next generation sequencing (NGS) has drastically changed the landscape of experimental biology, not only by helping to characterize cellular networks to an unprecedented level, but also by generating vast quantities of data. Dozens of tools currently exist for Illumina FASTQ file analysis, however, as big data analysis becomes an increasingly needed skill in biology, the demand for versatile user-friendly applications also rises. As NGS becomes simpler and widespread, so must its respective data processing. Therefore, there is a need for intuitive, reproducible, and versatile tools that can handle the sometimes overwhelming initial raw data processing step.

NGS applications often require features to be extracted and counted from FASTQ files for downstream analysis. For example, several analysis tools and scripts exist for systematic reverse genetic screens, such as CRISPRi-Seq [1] and random-barcode sequencing (RB-Seq) [2, 3]. However, at the moment, such pipelines are either overspecialized, complex, or requiring informatics skills beyond the average user [4-6]. Furthermore, pipelines often process NGS data as part of a downstream data analysis setup (i.e. differential read count). Depending on the experiment, it might be beneficial to uncouple one from the other. Current approaches are also limiting in throughput in regards to searching and returning reads with specific sequences, especially when considering sequence mismatches, and nucleotide wise Phred score filtering. This is particularly important in the cases where a user wants to control several processing parameters to fit their experimental setup.

Here, we introduce 2FAST2Q, a fast and versatile FASTQ file processor for extracting and counting sequence occurrences from raw reads. 2FAST2Q requires no installation, and works in all common operative systems. As a proof of concept, we show that 2FAST2Q efficiently and reliably counts single guide RNA (sgRNA) features in FASTQ files originating from published prokaryotic and eukaryotic CRISPRi-Seq experiments, and that it can be used for any *de novo* sequence searching, or for extracting any kind of sequences from FASTQ files.

## Results

### Developing 2FAST2Q

Typical NGS data generated by illumina sequencing is delivered in the form of a so-called FASTQ file. A FASTQ file is a large text file that contains the read DNA sequences with their respective quality scores, typically existing in its compressed form with the extension *.fastq.gz. A major goal when doing targeted (amplicon) sequencing is to know the abundance of each target within the sample. To that end, we wrote the Python-based tool called 2FAST2Q (Fig. 1). 2FAST2Q is able to efficiently extract, align, filter, and count DNA sequences from standard FASTQ files in a single step. 2FAST2Q also performs mismatch sequence searching, nucleotide Phred score quality filtering, and handles .gz files. The program exists as an easy to use intuitive executable version for MS Windows, macOS, and Linux, requiring no installation. Alternatively, 2FAST2Q is also available as a Python3 package in the PyPI repository, and can be installed with the “pip install fast2q” command. As input, 2FAST2Q requires only a FASTQ /gz file, and, when reference feature sequences exist (i.e.: sgRNAs, barcodes), a .csv file with all the lookup DNA sequences. As an output, 2FAST2Q returns an ordered .csv file with all the raw feature counts per condition, as well as quality control statistics (Fig. 1B-C). 2FAST2Q contrasts with other current methods by being easy to setup and intuitive to use (Fig. 1A), while simultaneously maintaining advanced configuration settings such as efficient mismatched sequence searching, and quality filtering.

**Figure 1.**
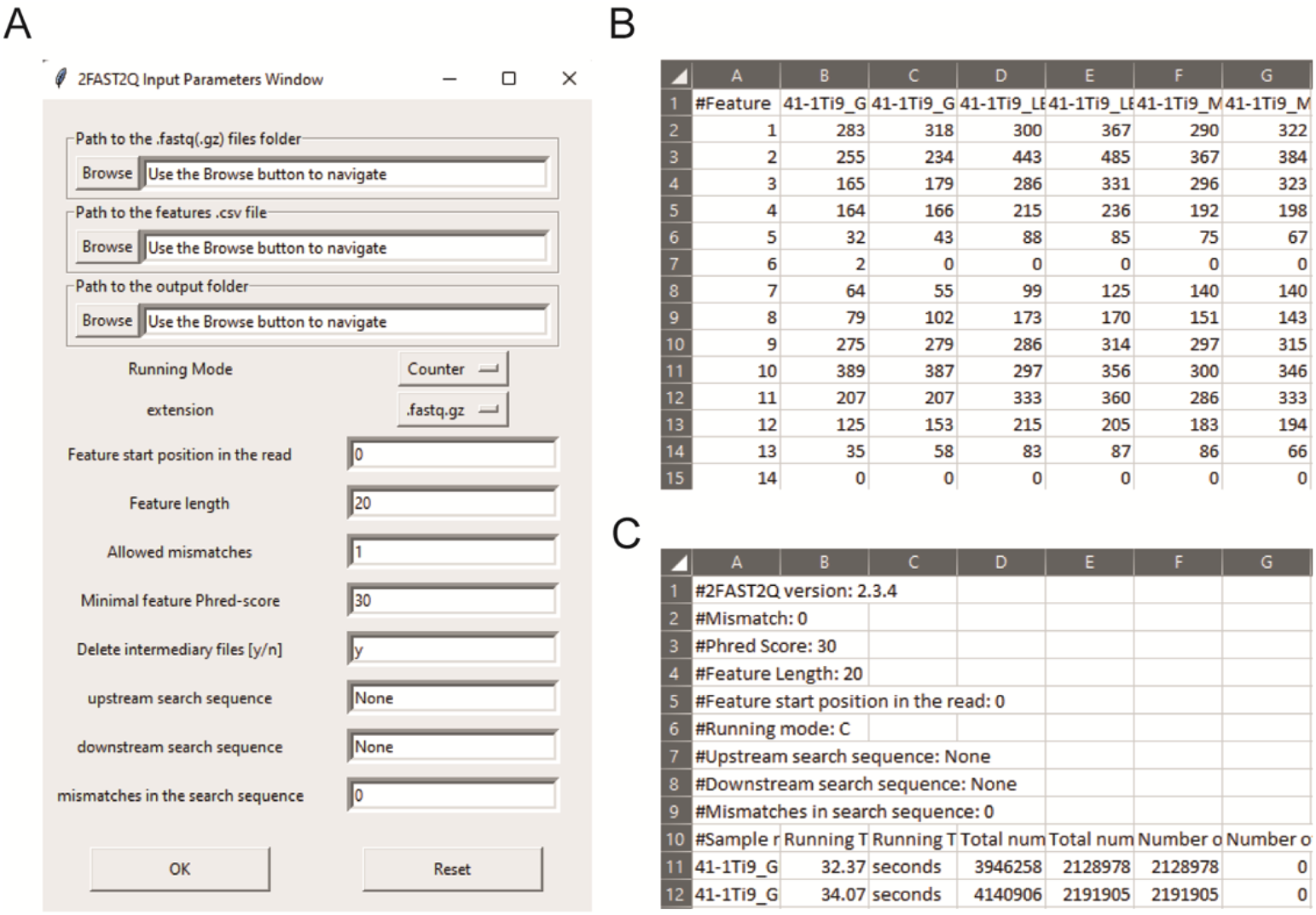
2FAST2Q interface, and outputs. (A) All program parameters are given by interacting with 2FAST2Q user interface. 2FAST2Q outputs two .csv files; a raw read count file for all samples (B), and a file with each sample statistics (C). Each independent file is considered to be a sample, and the file name the sample name.

### Counting features using 2FAST2Q

An important feature of performing CRISPRi-seq or RB-seq is to obtain reliable counts of each single guide RNA (sgRNA) or barcode, for any experimental condition. When using 2FAST2Q in “counting mode”, (i.e.: for CRISPRi-Seq, or sequence barcode counting), it can be used to quickly obtain an absolute feature sequence count from FASTQ files. When in “extract and count mode”, the program doesn’t require the input of any feature sequences, and will retrieve the count of all found read sequences. In both cases, the program can search for any feature by either specifying a starting read position, or by providing upstream and/or downstream constant search sequences. The feature length must be specified, except in the latter (Fig. 2).

**Figure 2.**
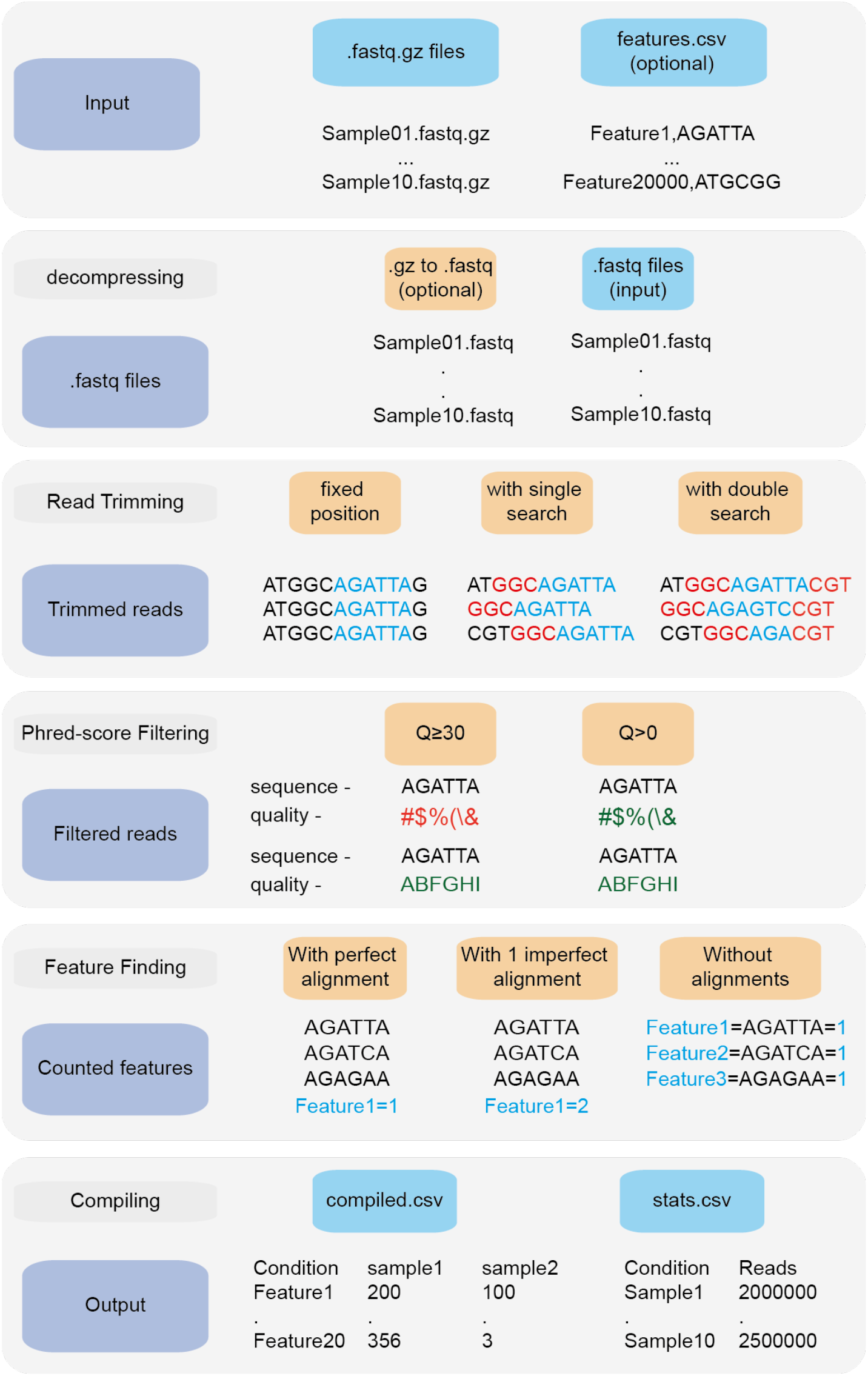
2FAST2Q pipeline. 2FAST2Q requires only .fastq.gz (or .fastq) files as input. When in alignment mode, a csv file with all the features must also be provided. 2FAST2Q performs all described steps automatically and without requiring external software. Trimming parameters, filtering scores, and mismatch tolerances can be easily adjusted using 2FAST2Q graphic interface.

### Benchmarking 2FAST2Q

2FAST2Q was initially benchmarked against a published CRISPRi-Seq dataset comprising 479M reads dispersed over 118 FASTQ files [7]. In this study, Rousset *et al* examined which genes are essential in *Escherichia coli* under different environmental conditions using CRISPRi-seq [7]. When only considering perfect alignments between a feature and a read, 2FAST2Q was able to output the final compiled sgRNA count table in 7 min on a personal desktop computer (33s per sample distributed over 10 parallel cores). When allowing for 1 mismatch in the sgRNA search count, the total run time only increased by 2min, to 9min. Under these program conditions, this corresponds to a more than 40x speed improvement over the use of similar purpose standard search functions, such as the Python regex module match function [8].

Using the same dataset published by Rousset, F. *et al*. [7], we assessed the impact of different initial 2FAST2Q parameters on both absolute feature counts, and on downstream data analysis. When not using any Phred-score filtering (Q≥0), and not allowing for any mismatches, we were able to fully recapitulate the reported total read counts/sgRNA for all conditions (Fig. 3E) (supplementary tables 1 and 2). However, high-quality read length has been reported to improve Illumina sequencing results interpretation [9]. We therefore implemented a filtering for nucleotide wise Phred-scores (Q), where all the sequenced nucleotide scores corresponding to the found feature read location are required to be above an indicated threshold. As expected, filtering using Q≥30, indicating a 0.1% probability of a nucleotide sequencing mistake, lowers the amount of reads/sgRNA. In some cases, by more than 1 order of magnitude (Fig. 3G). However, when considering the millions of reads generated by a typical sequencing experiment, the presence of mismatches in high quality reads is a likely event (any length of 20 nucleotides with Q≥30 have, at most, a 2% chance of having a mismatch: 0.001 * 20 = 0.02). We therefore implemented feature mismatch search where a read is considered valid if it unambiguously aligns to a single feature for any number of considered mismatches, thus retrieving more high quality reads, especially from lower overall quality sequencing runs. Allowing for mismatches expectedly increased the number of reads/feature (Fig. 3A, 3C, 3I and 4A), without sacrificing total run time (Fig 4B) (supplementary tables 3 and 4). As an extreme benchmark case, we allowed for the same number of mismatches as the feature length (20bp) (Fig. 3I-J). In practice, these parameters mapped any read to its closest feature, meaning the sequence that unambiguously differs the least from the read. This is performed by an inbuilt safety mechanism, where if more than one feature possible matches the read at the lowest amount of allowed mismatches (i.e. 1), the read is always discarded, but otherwise kept. In regards to the Rousset, F. *et al*. dataset, which is of high quality, these parameters recovered on average 3% more reads/sgRNA (Fig 4A). However, it is conceivable that the use and outcome of these parameters varies depending on the experimental setup, requiring careful consideration before proceeding to downstream data analysis.

**Figure 3.**
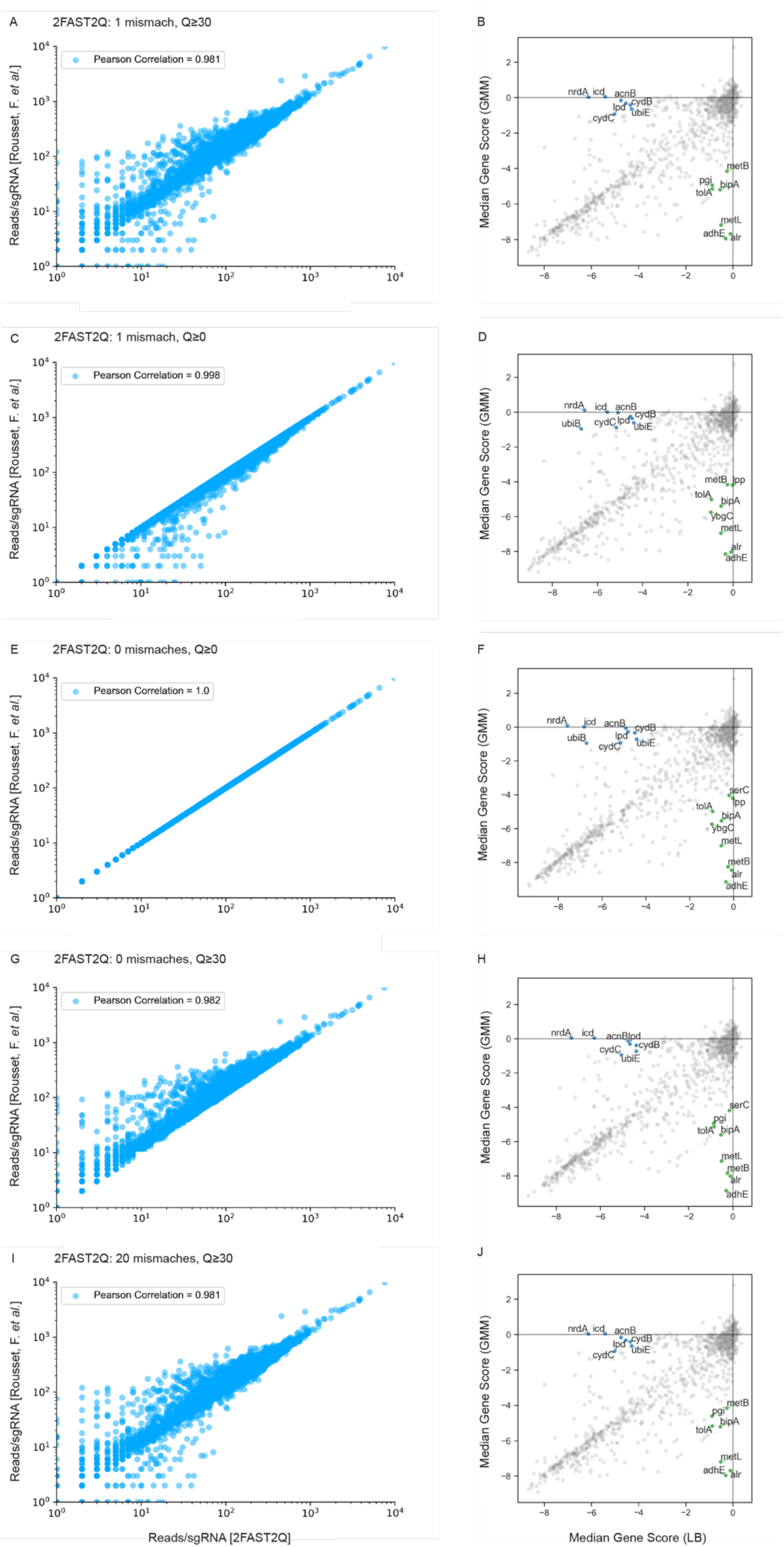
Absolute read counts/sgRNA for the Rousset, F. *et al*. dataset MG1655 LB 1 condition [7]. The total read counts using different 2FAST2Q mismatch and/or quality filtering inputs are plotted against those reported by Rousset, F. *et al*. Pearson correlation for each plot is also shown. Plots B, D, F, H and J were generated using an adaptation of the published Jupyter notebook analysis pipeline, and highlight the significant genes (absolute fold change ≥ 4 in one condition and ≤ 1 in the other) when using different 2FAST2Q input parameters. green: fold change < 4 in GMM media, and > -1 in LB; blue: fold change < 4 in LB media, and > -1 in gut microbiota medium (GMM);

**Figure 4.**
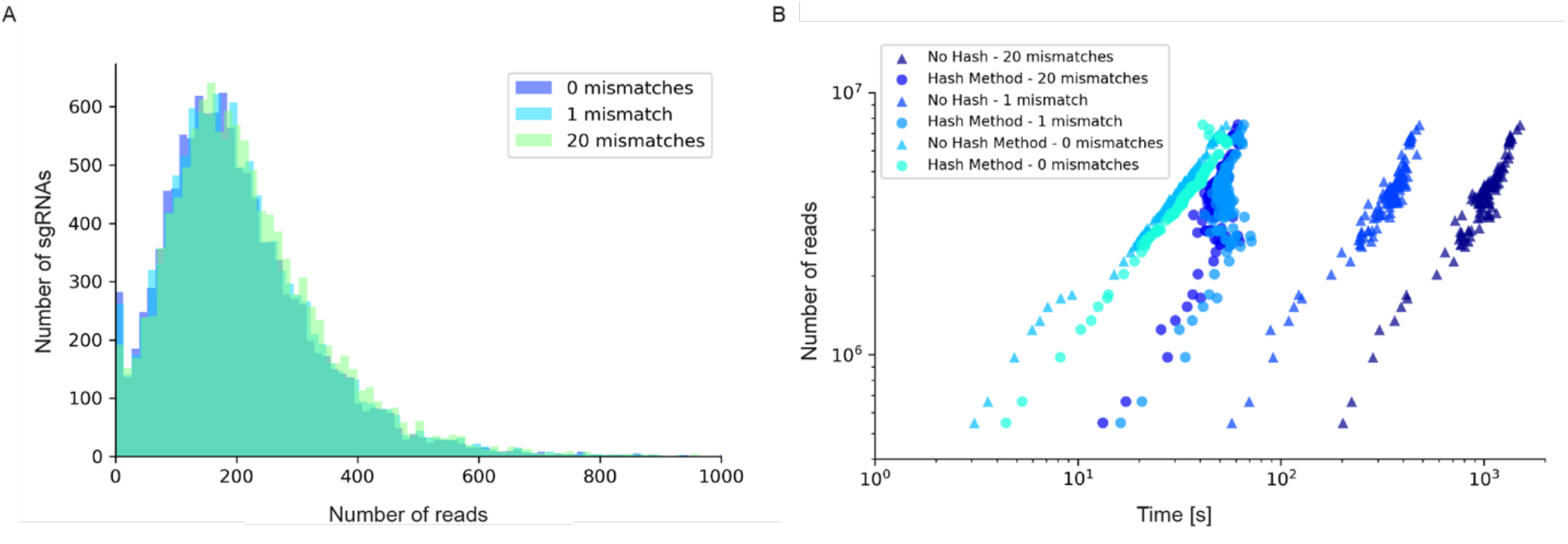
Read/sgRNA distribution and runtime analysis of 2FAST2Q with different mismatch parameters and algorithms. Data analysis was performed on the Rousset *et al*. “UTI89_T0” fastq sample [7] when submitted to 2FAST2Q analysis with either 0, 1, or 20 mismatches (and Phred score ≥ 30). Increasing mismatches allows for greater read recovery by matching a given read to its closest matching (and thus most likely) feature. A) The median reads/sgRNA increased from 182 to 187, and then to 196, when considering 0,1, and 20 mismatches, respectively. B) 2FAST2Q runtime analysis demonstrates the efficiency of real time creation of pre-processed failed/passed read hash tables (see methods) vs. the “no hash” method, where each read is always processed *de novo* for mismatches.

### Higher stringency parameters can aid in biological discovery

We used the Jupyter notebook analysis pipeline published by Rousset, F. *et al*. [7] to assay how these different read processing scenarios impact downstream analysis. Using the different read count tables directly outputted by 2FAST2Q, we calculated and compared the median gene scores as defined by Rousset *et al*. (essentially, the median of the log_2_ fold change for each feature in all experimental replicates) for the LB medium and gut microbiota medium (GMM) conditions. Using more stringent criteria than Rousset, F. *et al*. (a gene is considered significant if it has an absolute gene fold change ≥ 4, instead of ≥ 3.5), we compared how different Phred-scores and mismatch filtering criteria influenced downstream analysis, namely how these criteria influence gene score calculations, and thus gene essentiality (Fig. 3B, D, F, H, J).

We observed a higher stringency for the 2FAST2Q parameters of 1 mismatch, and base pair quality filtering of ≥ 30 (Fig 3B), with fewer genes being considered essential for any given condition with these criteria than with the criteria that recapitulate the published data (0 mismatches allowed, and no Phred-score consideration) (Fig. 3F). As expected, different read filtering criteria resulted in fold change differences, and consequently in differences in the genes considered essential for these conditions. What criteria to use would depend on the specifics of each individual experiment. The default 2FAST2Q parameters uses a Phred-score of ≥ 30, while allowing up to 1 mismatch (representing for any 20 basepair (bp) sequence, a 5% bp deviation error with, at maximum, a 2% chance of any nucleotide being wrongly sequenced). As shown in Fig. 3, although the default setting of 2FAST2Q give slightly fewer significant hits, they were all also reported by Rousset *et al*. It is also conceivable for a user to be interested in aligning all reads to their closest matching feature. This is possible by setting the total amount of mismatches to the same length of the feature. In this case, the 2FAST2Q inbuilt alignment safety mechanism will prevent ambiguous read alignments from being considered.

### 2FAST2Q dynamically performs FASTQ feature extraction

Under certain experimental setups, the extraction of features from FASTQ files might require the use of a dynamic trimming and search function (i.e.: when the location and/or size of the feature differs from read to read) (Fig 2). In this case, a delimiting search sequence of any length (up and/or downstream of the feature) can be provided. Similar to feature mismatch search, an arbitrary number of mismatches can also be indicated for the search sequence-based trimming. 2FAST2Q will then efficiently search each read for the indicated sequences, returning the correctly trimmed read for further processing. As a proof of concept, we used a published CRISPRi-Seq dataset by Wei *et al*. [10], where dynamic read trimming was required. In this study, a CRISPRi screen was performed using Vero-E6 cells (kidney epithelial cells from an African green monkey) infected with SARS-CoV-2 to identify host genes important for viral replication [10]. Dynamic trimming allows each read to be independently trimmed based on the relative location of the found search sequences, thus always returning the correct feature location. Using this method, we submitted 6 FASTQ files (SRR14668185 - SRR14668190) to 2FAST2Q processing. As search sequence we used a 10bp upstream constant sequence (CGAAACACCG), allowing for 1 mismatch search error in this sequence. We used the provided list of 84,953 sgRNA sequence features, and ran 2FAST2Q (Q≥30, 0mismatches). 2FAST2Q simultaneously processed all 6 samples, comprising 324M reads, within 8 minutes on a standard desktop PC (supplementary table 5 and 6). This result corresponds to a slowdown of only 22% (speed comparisons were determined using processed reads/second) when compared with the non-dynamic feature extraction process, such as the one we used for the same parameter 2FAST2Q run with the Rousset, F. *et al* dataset [7].

Recently, Bosch *et al*. published a CRISPRi-Seq experimental setup with variable length sgRNAs [11]. In this case, 2FAST2Q was also able to extract, count, and align all the found features in a *Mycobacterium tuberculosis* dataset (SRR13734827), to the provided 96,700 long sgRNA file, albeit using 2 delimiting constant search sequences (upstream: GTACAAAAAC; downstream: TCCCAGATTA), while allowing for 1 mismatch in each. The returned variable length sequence between the two constant search sequences was used for perfect match alignment against the sgRNAs (supplementary table 7 and 8). When compared with the non-dynamic extraction process, a slowdown of 44% was observed, in line with what was observed for the Wei *et al*. dataset.

As the sequence search algorithm uses a similar process to the one used for feature alignment mismatch, a similar speed improvement over standard Python functions is also obtained. Together, these benchmarks demonstrate that 2FAST2Q is a versatile and quick computational tool that can extract relevant features and counts from FASTQ files.

## Discussion

We developed a fast, intuitive, and easy to use tool for counting sequence occurrences in FASTQ files. We have recently implemented the use of 2FAST2Q in our CRISPRi-seq pipeline and have found it very useful in the first step of data analysis [12]. Due to the parameter versatility of 2FAST2Q, the program covers most current user applications that require the extraction and counting of specific feature sequences, such as CRISPRi-Seq and RB-Seq. Despite only handling single-ended FASTQ files at the moment, the processing of paired-ended files is possible by running two separate instances of 2FAST2Q. The program will automatically compile all samples at the end if all intermediary files of the first run are copied to the output folder of the second instance while processing.

Current methods require users to handle several different software pipelines in order to extract relevant data from FASTQ files. 2FAST2Q is a standalone program that can, in a single step, efficiently and quickly perform nucleotide wise quality filtering, mismatch sequence searching, *de novo* feature extraction, and sequence occurrence counting. 2FAST2Q outputs an individually compiled, easy to interpret, excel readable.csv file with all the feature counts per sample, alongside a file with relevant sample statistics.

2FAST2Q fully recapitulated the feature counts reported by Rousset, F. *et al*. for all conditions when using the same filtering criteria. However, upon analyzing the published CRISPRi-Seq dataset using a Phred-score filtering of ≥30 and allowing, at most, 1 mismatch between read and feature, fewer GMM condition specific genes with a fitness defect were found (Fig 3F and B), indicating the higher stringency of these parameters. It will be interesting to see whether the hits identified under these more stringent, high-quality filtering and selection conditions results in fewer false positives after following up by experimental validation. In any case, we implemented this option as the default mode as it reduces raw data noise while maximizing read recovery. 2FAST2Q was also successful at extracting features starting at different positions per read when using a published dataset of a CRISPRi screen on eukaryotic cells that were infected with SARS-CoV-2 [10]. 2FAST2Q inbuilt search functions also allow for more complex experimental setups. For example, recent work by Bosch *et. al* applied CRISPRi-Seq with variable length sgRNAs to identify conditionally essential genes in *M. tuberculosis* [11]. By providing up and downstream search sequences, 2FAST2Q efficiently, in a single step, extracted these sgRNAs. In the case of experiments with more than one feature per read, such as with dual barcode sequencing, or dual CRISPRi-Seq, it is conceivable that 2FAST2Q could also be used, taking into account that the parameters need to be adjusted to capture different features per read each time, and by compiling the data at the end.

Besides being able to align and count provided features in FASTQ files, 2FAST2Q is also able to extract and count all unique read sequences when in “extract and count mode”. In this case, all different sequences that fulfill the required parameters are returned, with any possible mismatches being accounted as distinct sequences.

As experiments that produce large datasets become more widespread, the need for versatile, fast and easy to use software that handles raw data becomes more pressing. It is thus our hope that 2FAST2Q can contribute to facilitate the processing of the large amounts of sequencing data originating from NGS studies.

## Methods

### Installation and code availability

All 2FAST2Q executable files can be downloaded from zenodo: https://zenodo.org/record/5410822. The code, usage instructions, and test datasets are available on GitHub: https://github.com/veeninglab/2FAST2Q. 2FAST2Q is also a Python package, and can be accessed on PyPI: https://pypi.org/project/fast2q/. When using the executable version on MS Windows or MacOS, no further installation is required and a double click on the executable should suffice. For a more in depth description, please see the online tutorial on https://veeninglab.com/2fast2q. 2FAST2Q is fully implemented in Python3.

### Usage considerations

All indicated 2FAST2Q running times were performed on a desktop PC with a 12 core 3.7GHz processor, and 32GB of RAM. However, 2FAST2Q runs on any up-to-date desktop or laptop. When using 2FAST2Q without mismatch search (perfect alignment only), sample processing should be in the order of seconds or minutes (after file decompression). When using the mismatch search on large datasets, it is possible for 2FAST2Q analysis to take several minutes per sample. When processing more than one sample, 2FAST2Q will automatically parallelize all analyses by distributing each sample per available processor core.

2FAST2Q fast sequence mismatch search function was possible due to the use of Python numpy [13] and numba [14] modules. An advanced and in-depth tutorial on 2FAST2Q parameters is available on GitHub and PyPI.

### 2FAST2Q algorithm

When initialized in standard feature count mode, 2FAST2Q will automatically decompress all FASTQ files, and create a hash table for all supplied sequence features. 2FAST2Q will then forward all samples to individual processes for parallel processing, which can be monitored via a progress bars (supplementary fig 1). Each FASTQ file is sequentially read, saving RAM space. Each loaded read is submitted to trimming based on the indicated parameters, either using a fixed position, or a dynamic search. The first assumes the presence of a fixed feature length in the same location for all reads. The second requires one or two search sequences. When one sequence (either up or downstream) is provided, 2FAST2Q will search the read until the sequence is found, and return the predetermined sized feature (again, either up or downstream). When two sequences are used, 2FAST2Q will return any feature within the found search sequences. The location and feature length parameter can thus be ignored in this latter scenario. A sequence mismatch search can also be performed. Following read trimming, the Phred-score corresponding to each nucleotide of the trimmed sequence is considered. If any of the scores is below the indicated parameter threshold, the read is discarded.

If the read passes quality control, alignment against the input features is finally attempted. Depending on the user input, any kind of feature alignment is performed using either mismatch search or not. By default, 2FAST2Q will always first check for a perfect match. Perfect matching uses hashing, directly comparing all features to the read sequence using hashing runtime complexity. When dealing with mismatches, 2FAST2Q will perform sequence search based on a faster custom made search algorithm. At first, all feature/search sequences are converted to their numerical binary form, subsequently reducing them to integer8 format using numpy. Sequence mismatches are counted by tracking the non-zero result positions of subtracting both sequences. As simple numpy constructs, arithmetic operations can be easily processed using the Python Numba module njit decorator. Therefore, all 2FAST2Q search functions are pre-compiled and effectively run at much faster speeds (supplementary figure 2). All read sequences searches, and features mismatch alignments are performed using this approach, allowing all search operations to run faster than standard Python code. Moreover, reads that fail to safely align, within the given parameters, to any of the provided features, are stored and used for quick hashed based comparison. The same is performed for reads that align with mismatches. By performing the much faster hashed comparison, this feature avoids the slower *de novo* mismatch search for previously seen same sequence reads. Runtime is thus decreased, paradoxically maintaining sample processing time as file size increase. “Already seen read” hashing is especially useful with large datasets comprising dozens of samples from the same sequencing run (Fig. 3B). In this case, the generated failed/passed read hash tables for each sample are compiled and used as a seed to the next batch of samples. Each new sample thus takes advantage of the already processed reads in a previous sample, avoiding reprocessing the exact same read several times.

A Python dictionary with a class feature count is used to keep track of all found aligned sequences. When no feature file is provided (i.e. when running in “Extractor+Counter” mode), all found read sequences are returned and counted. Each FASTQ file will originate a unique output file. At the end of the analysis, all samples files are compiled into a single file, which can be readily used for downstream applications.

## Acknowledgements

We thank Julien Dénéréaz and Vincent de Bakker for their software tests, and all members of the Veening lab for helpful discussions. Work in the Veening lab is supported by the Swiss National Science Foundation (SNSF) (project grants 310030_200792 and 310030_192517), SNSF JPIAMR grant (40AR40_185533), SNSF NCCR ‘AntiResist’ (51NF40_180541) and ERC consolidator grant 771534-PneumoCaTChER. AB is the recipient of the Fundação para a Ciência e Tecnologia (FCT) individual research fellowship (PD/BD/128006/2016).

## Supplementary information

**Supplementary figure 1.**
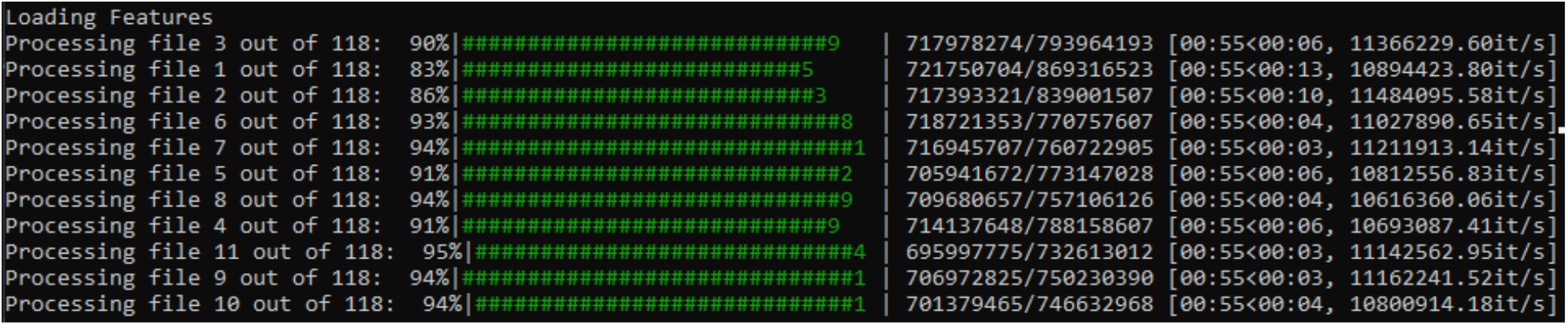
2FAST2Q parallel sample processing screen.

**Supplementary figure 2.**
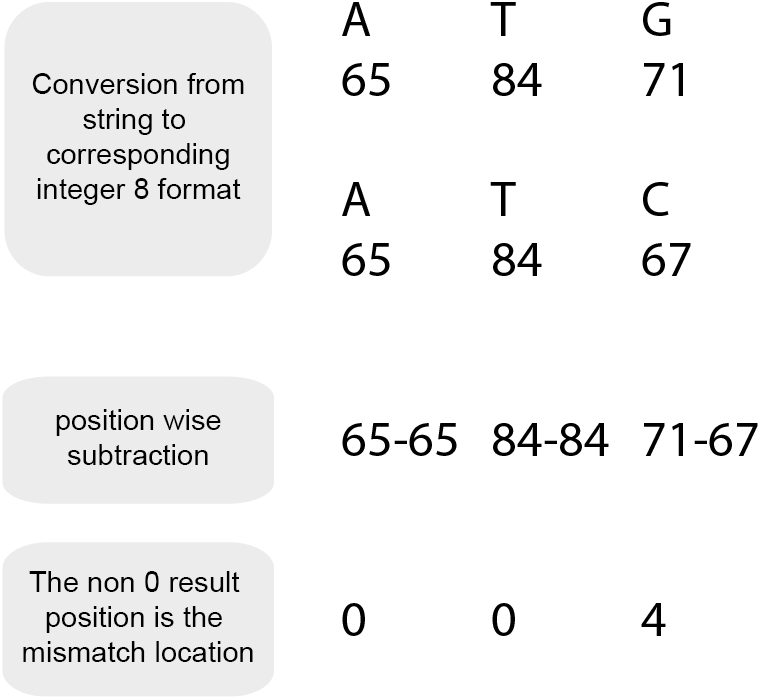
2FAST2Q mismatch search algorithm. When dealing with mismatches, 2FAST2Q will preemptively convert all the input feature sequences into their respective binary integer format using 8bits encoding. This step ensues faster downstream processing, decreases RAM usage, and allows mismatches to be calculated by a simple subtraction performed at machine code speed.

## Notes

https://veeninglab.com/2fast2q

### Competing Interest Statement

The authors have declared no competing interest.

